# Quantifying metacognitive thresholds using signal-detection theory

**DOI:** 10.1101/361543

**Authors:** M.T. Sherman, A.K. Seth, A.B Barrett

## Abstract

How sure are we about what we know? Confidence, measured via self-report, is often interpreted as a subjective probabilistic estimate on having made a correct judgement. The neurocognitive mechanisms underlying the construction of confidence and the information incorporated into these judgements are of increasing interest. Investigating these mechanisms requires principled and practically applicable measures of confidence and metacognition. Unfortunately, current measures of confidence are subject to distortions from decision biases and task performance. Motivated by a recent signal-detection theoretic behavioural measure of metacognitive sensitivity, known as meta-*ď*, here we present a quantitative behavioural measure of confidence that is invariant to decision bias and task performance. This measure, which we call *m*-*distance*, captures in a principled way the propensity to report decisions with high (or low) confidence. Computational simulations demonstrate the robustness of *m*-*distance* to decision bias and task performance, as well as its behaviour under conditions of high and low metacognitive sensitivity and under dual-channel and hierarchical models of metacognition. The introduction of the *m*-*distance* measure will enhance systematic quantitative studies of the behavioural expression and neurocognitive basis of subjective confidence.

## 1. Introduction

Effective decision-making often requires the evaluation of the accuracy or effectiveness of our decisions, a process usually referred to as metacognition (‘cognition about cognition’) (Flavell, 1979). For example, learning is facilitated when we are adept in retrospectively identifying our mistakes (Guggenmos, Wilbertz, Hebart, & Sterzer, 2016; Yeung & Summerfield, 2012). Similarly, group decision-making improves when people communicate how confident they are in their beliefs (Bahrami et al., 2010; Zarnoth & Sniezek, 1997). Confidence judgements are usually strongly correlated with the signal to which they pertain: confidence increases with both the strength and reliability of bottom-up (e.g., sensory) information (Baranski & Petrusic, 1998; Maniscalco & Lau, 2012; Peirce & Jastrow, 1884; Spence, Dux, & Arnold, 2015). However, confidence is also influenced by top-down processes such as attention (Kanai, Walsh, & Tseng, 2010), prior beliefs about, or experience with, the percept (Melloni, Schwiedrzik, Müller, Rodriguez, & Singer, 2011; Sherman, Seth, Barrett, & Kanai, 2015), as well as the subject’s previous confidence reports (Guggenmos et al., 2016; Rahnev, Koizumi, Mccurdy, Esposito, & Lau, 2015). The ways in which these different sources of information are combined into confidence judgements, the neurocognitive mechanisms underlying this integration, and even the identity of the information sources relevant to confidence all remain open questions.

Three quantities stand out as especially relevant in the study of metacognition and confidence: (i) the objective sensitivity to some signal of the participant during a task, referred to as type 1 sensitivity; (ii) the tendency to report with high confidence, known as metacognitive or type 2 threshold; and (iii) metacognitive or type 2 sensitivity, which captures the trial-by-trial correspondence between confidence and decision accuracy. While these quantities are conceptually distinct, they tend to correlate in empirical settings. For example, greater objective sensitivity is often associated with higher average confidence (Grimaldi, Lau, & Basso, 2015), which makes sense if type 1 and 2 decisions are constructed from similar bottom-up information (Fetsch, Kiani, Newsome, & Shadlen, 2014; Kvam & Pleskac, 2016). The existence of such empirical correlations raise the question of whether they reflect features of underlying neurocognitive mechanisms, or whether they arise because of the measures used to operationalise decision making behaviour. Advancing our understanding of metacognition and confidence therefore requires measures that are sensitive to, and not confounded by, these conceptual distinctions.

A powerful framework for developing such measures is provided by signal detection theory (SDT), which affords a principled way of decoupling sensitivity on a task from thresholds for both objective (e.g. “Left vs. Right”) and subjective (e.g. “Confident” vs. “Guess”) reports (Green & Swets, 1966). For binary type 1 (i.e. objective) decisions, SDT assumes that the brain represents distinct likelihood distributions over the signal representation for each option. A decision threshold determines whether a particular representation will be classified as having been driven by one or the other signal class. The separation of the two distributions represents the subject’s type 1 sensitivity: greater separation indicates a greater ability to distinguish the two types of signal. Thus, modelling subject’s responses using type 1 SDT enables computation of (theoretically) independent measures of task performance and decision bias, known as *ď* and *c* respectively (see fig 1C, top panel).

**Figure 1.**
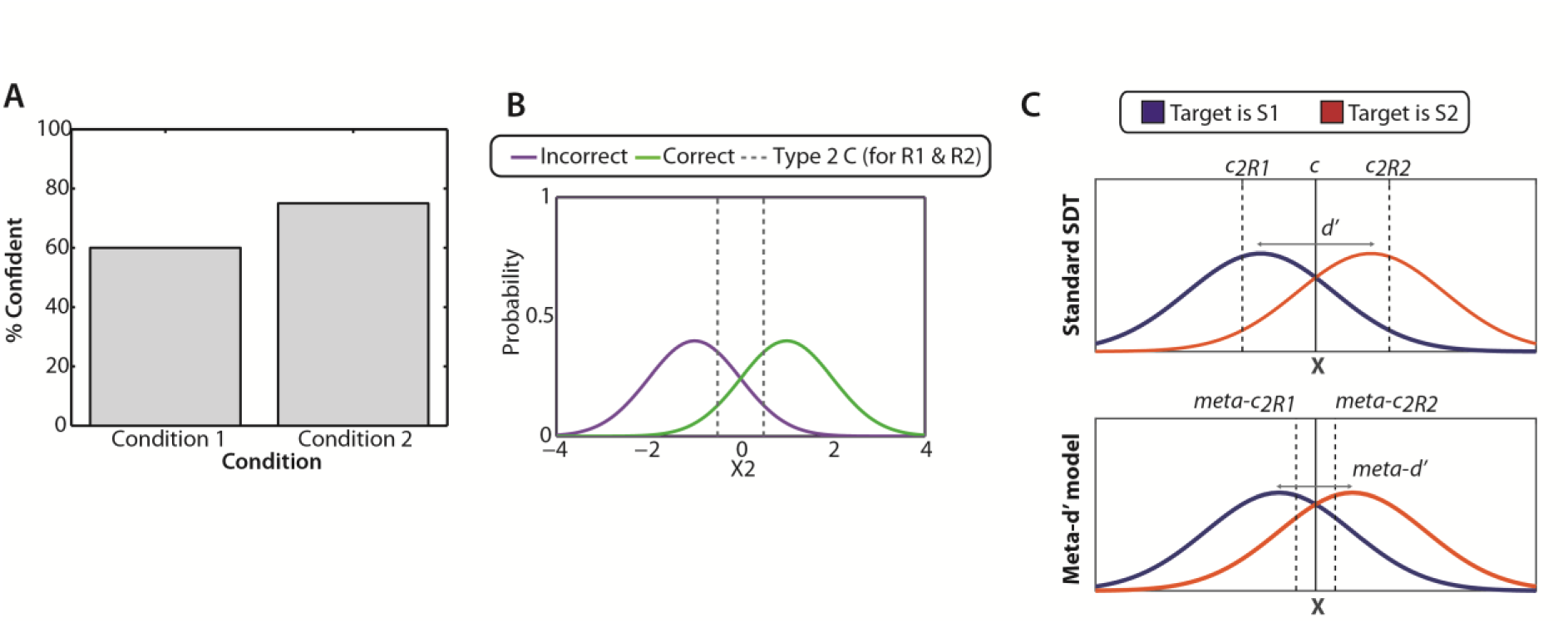
Different measures of metacognitive threshold. **A.** Mean confidence as a function of condition (or group, response type etc.) **B.** Type 2 SDT (Evans & Azzopardi, 2007) used to compute type 2 C, which reflects the observer’s bias towards reporting confidence. Unfortunately, the theoretical assumptions needed to justify type 2 C do not hold (Galvin et al., 2003). **C, top.** Standard SDT. This assumes that an observer’s stimulus sensitivity (denoted ď) is determined by the separation between two Gaussian likelihood functions, P( X | S_1_) in blue and P( X | S_2_) in red. The observer’s decision threshold (black solid line), denoted c, determines the evidence required to report a decision as R_1_ (which we call a negative decision) or R_2_ (which we call a positive decision). Here decisions will be unbiased, because c is located where two likelihood functions cross; other locations of c lead to biased decisions. Flanking confidence criteria (dashed lines) for negative and positive decisions are denoted c2_R1_ and c2_R2_ respectively. **C, bottom.** The meta-ď model. This is the model that would be used by an SDT-ideal observer with the same type 1 decision bias, and same type 2 response data as the observer. The variables have the same interpretation as those in the standard SDT model, but the values ď, c2_R1_ and c2_R2_ take can differ if the observer’s metacognitive sensitivity departs from SDT-ideal predictions. Accordingly, each parameter is prefixed by ‘meta-’ (Maniscalco & Lau, 2012).

Recent developments in SDT have also led to principled methods for quantifying metacognitive sensitivity such that the estimate is independent of type 1 performance and both type 1 and 2 decision biases, called meta-*ď* (Barrett, Dienes, & Seth, 2013; Maniscalco & Lau, 2012). This allows, for example, to demonstrate changes in meta-memory during development without the confound of increasing recognition performance (Fandakova et al., 2017).

While meta-*ď* is now well-established as a principled measure of metacognitive sensitivity, less attention has been paid to how this approach can be used to quantify people’s metacognitive *threshold*, which is of independent interest as a property of metacognition. Metacognitive sensitivity quantifies the extent to which confidence judgements discriminate accuracy. Metacognitive threshold is the amount of additional evidence to make a report with confidence. Just as decision bias and decision accuracy provide complementary information about objective performance at the type 1 level, metacognitive threshold and metacognitive sensitivity provide complementary information about metacognitive (type 2) performance.

Thus far, most empirical work examining metacognitive thresholds has used one of two measures: the proportion of confident responses made (figure 1A); or the measure of metacognitive bias type 2 C (figure 1B). The measure type 2 C (Kunimoto, Miller, & Pashler, 2001) is a direct analogue of type 1 c in standard SDT, but is constructed by grouping trials according to confidence and accuracy instead of by signal class and response.

Unfortunately, it has been demonstrated that despite the conceptual analogy, type 2 C is not independent of metacognitive sensitivity (Galvin, Podd, Drga, & Whitmore, 2003). Therefore, the need for a measure of metacognitive threshold that is independent of (type 1) response bias remains. Without such a measure, we cannot be sure that changes to confidence reports (as measured by proportion of confident trials, type 2 C or otherwise) are driven by changes in metacognitive decision making, rather than driven by changes at the type 1 (objective) level, i.e. decision biases. Addressing this need, we develop and illustrate an alternative quantification of metacognitive threshold based on the computation of meta-*ď*, which is approximately (theoretically) invariant to the confounds of decision bias and sensitivity (Barrett et al., 2013).

First we outline signal detection theory and summarize how meta-*ď* is estimated. Next, we propose a formal and principled measure of metacognitive threshold (termed ‘*m*-*distance*’), based on *meta*-*ď*, and finally, we simulate the behaviour of this measure to show that it adequately captures the intuitive description of a metacognitive threshold and is robust to changes in detection sensitivity, decision criteria, and differing degrees of metacognitive sensitivity. We compare its performance under different conditions with that of raw confidence, and finally, demonstrate the robustness of the measure under differing trial counts, concluding that the measure performs best when the experimenter collects at least 300 trials.

## 2. Quantifying confidence in Signal Detection Theory

### 2.1 Signal detection theory

Signal detection theory (Green & Swets, 1966; Macmillan & Creelman, 2004) models binary judgements as a process by which the observer categorises an internal response X to a stimulus S_1_ or S_2_ as having arisen from one of two respective Gaussian generative likelihood distributions P(X|S_1_) or P(X|S_2_). The separation between the central tendencies of these distributions reflects the observer’s sensitivity to the task (in the standard deviation of P(X|S_1_) units). A larger separation between the two Gaussians indicates greater type 1 (i.e. objective) sensitivity, because the observer generally has less uncertainty about which stimulus would have generated the sensory evidence observed. The separation is denoted *ď* (see figure 1C, top panel).

The response R made by the observer is determined according to a decision threshold or criterion θ such that:

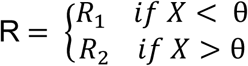

Several interpretations and formalisations of θ exist in the literature, and in the present paper we use the measure *c.* This quantity reflects the distance from the intersection of the two likelihood functions. When *c* is equal to zero the subject is unbiased, whereas *c* less than or greater than zero indicates a bias towards reporting R_2_ or R_1_ respectively. Importantly, if the likelihood functions have equal variances (as is usually assumed on two alternative force-choice discrimination tasks, but not in general for detection tasks) then *c* and *ď* can be unambiguously determined from the hit rate (proportion of *R*_2_ reports when the stimulus is *S*_2_) and false alarm rate (proportion of *R*_2_ reports when the stimulus is *S*_1_):

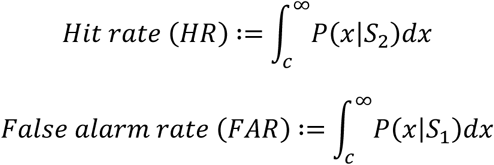

and

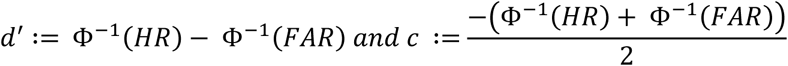

where Φ^−1^ is the inverse of the cumulative density function of the standard normal distribution of mean 0 and standard deviation 1 (also known as the Z-function).

### 2.2 Modelling confidence in SDT

To accommodate confidence judgements in SDT two (or more) flanking confidence criteria can be placed around the decision criterion *c*, denoted *c2*_R1_ and *c2*_R2_ (figure 1C, top panel). When an R_1_ response is made, a confident R_1_ response requires the evidence to also have surpassed the *c2*_R1_ threshold. Analogously, an R_2_ response is confident if the evidence X is less than *c2*_R2_. The decision rule for the confidence judgement C is thus formally

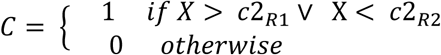

These two thresholds must be well-ordered, so that *c2*_*R*1_ ≤ *c2*_*R*2_.

Estimating confidence criteria when they are placed over the type 1 decision axis is straightforward. First, ‘confident R_1_’ responses are re-classed as ‘R_1_’ and the others are re-classed as ‘R_2_’. Calculating *c* on these reclassified trials gives *c*2_R1_. The same procedure is used to estimate *c2*_R2_ except that only ‘confident R_2_’ trials are classed as ‘R_2_’ and the rest are classed as ‘R_1_’. This procedure is illustrated in Table 1.

**Table 1.**
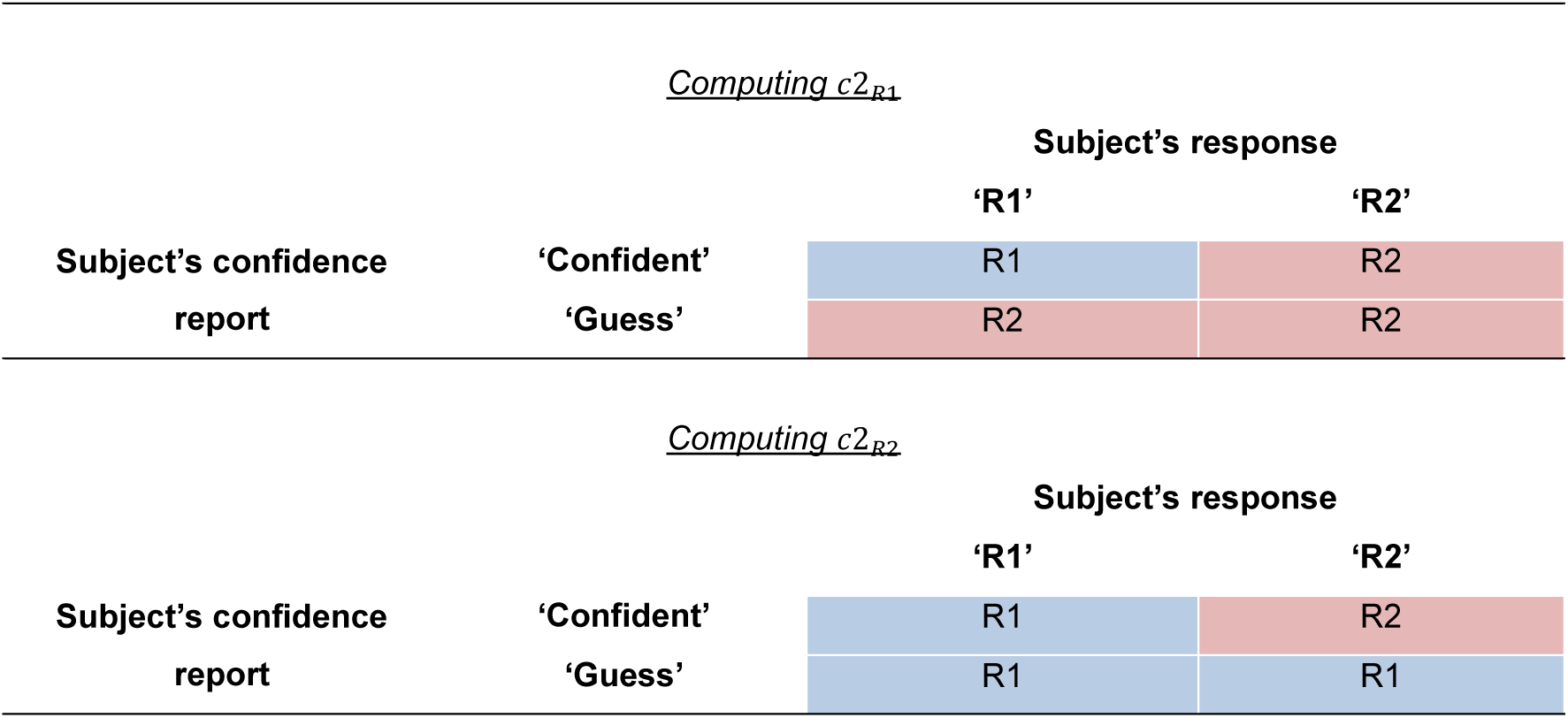
Reclassification of responses into the R1 (blue cells) or R2 (red cells) classes, according to a subject’s reported judgement and confidence rating.

This reclassification method assumes that human observers use the same evidence distributions to construct their objective and subjective judgements, reflecting so-called single-channel or ‘read-out’ models of confidence (Maniscalco & Lau, 2016). However, single-channel models struggle to accommodate dissociations between confidence and accuracy. For example, confidence is less sensitive to signal variance than the objective judgement (Zylberberg, Roelfsema, & Sigman, 2014). This strict (and as yet un-validated) model assumption renders *c2*_R1_ and *c2*_R2_ a poor choice for representing confidence in SDT.

### 2.3 Quantifying metacognition using meta-*ď*

As mentioned, quantifying confidence as *c2*_R1_ and *c2*_R2_ necessitates that the experimenter makes the strong assumption that confidence judgements are constructed from the same signal representation as the type 1 judgement. The measure *meta*-*ď* quantifies metacognitive sensitivity without having to make this strict assumption, and accordingly it is increasingly accepted as the gold-standard measure of metacognitive sensitivity, and has been successfully applied in a number of contexts (e.g. Allen et al., 2016; Fleming et al., 2015). Essentially, *meta*-*ď* is the type 1 ď that could have led to the observed type 2 responses, if the observer had used the exact same evidence available for making both type 1 and 2 decisions. The subject’s metacognitive *efficiency* is then quantified as the discrepancy between the so obtained meta-ď measurement and the actual type 1 ď. Importantly, their ratio (*m*-*ratio* ≡ *meta*-*ď*/*ď*) is therefore a meaningful measure because they are in the same (*ď*) units.

In practice *meta*-*ď* is estimated by identifying for a given type 1 *c*, the type 2 ROC curve, and from this the SDT model (i.e. the parameters *ď* and *c2*) providing the best fit to the observed confidence and accuracy data. It is assumed that all evidence available for the objective (R) judgement is available for the subjective (C) judgement. The parameters of the *meta*-*ď* model are prefixed with ‘meta-’ to distinguish them from the ‘type 1’ model acquired from the empirical data (see Fig. 1C, bottom panel). Because the SDT-optimal observer uses all of the available evidence in their confidence judgements, confidence thresholds meta-*c2*_Ri_ and meta-*c2*_R2_ are placed over the type 1 decision axis and are left free to vary. The value of *ď*, denoted meta-*ď*, is also left free to vary. The value of *c*, denoted meta-*c*, is fixed such that meta-*c*/meta-*ď* = *c*/*ď*. This means that the hypothetical optimal (‘meta’) observer will make the same proportion of “yes” and “no” responses as the subject, i.e. they have the same type 1 decision bias.

There are several methods for estimating the ‘meta’ quantities: one is analytic (Barrett et al., 2013), obtained as a weighted mean of exact solutions for equations for R1 responses and R2 responses; one is based on either maximum likelihood estimation or sum of squares minimization (Maniscalco & Lau, 2012) for the combined set of equations for both R1 and R2 responses; a third method involves estimating group-level parameters by using hierarchical Bayesian estimation to incorporate uncertainty about single-subject parameter estimates (Fleming, 2017).

As mentioned, the value of meta-*ď* obtained can be compared with the measured *ď* to give the efficiency of the observer’s metacognitive judgements: when meta-*ď* = *ď* the observer is using all of the available information for their objective judgement in their confidence judgement, and the two SDT models (from which *ď* and meta-*ď* arise) are the same; when meta-*ď* < *ď* there is signal loss between the two responses, and the confidence-accuracy correspondence exhibited by the subject can be achieved with lower sensitivity by an SDT-optimal observer. In other words, under conditions of greater uncertainty about the cause of sensory signals (i.e. lower *ď*) the SDT-optimal observer will achieve the same confidence-accuracy correspondence as the subject. If meta-*ď* > *ď* then the inference is that the subject is ‘super-optimal’, that is, performs better than the SDT-ideal observer. This might reflect further evidence accumulation between the objective and subjective judgement (and violation of the feed-forward assumptions of SDT), among other possibilities (for example, the contribution of additional signal sources).

### 2.4 Using the meta-*ď* computation to estimate metacognitive threshold

We next address how the framework of *meta*-*ď* can be extended to quantify metacognitive threshold. As mentioned in the Introduction, a principled measure of metacognitive threshold should reflect the additional evidence required for a decision to be made with high confidence. Accordingly, it should be sensitive only to the difference between type 1 and 2 criteria, not to their absolute positions on any scale. It should also be independent of metacognitive sensitivity *meta*-*ď*, just as *meta*-*ď* is invariant under changes to decision and confidence criteria.

The additional evidence required for a decision to be reported with high confidence is reflected in the distance between the decision threshold *c* and each confidence criterion, which we denote *m*-*distance*, or *m*-*distance_R1_* and *m*-*distance_R2_* when considering them for each response class separately. We cannot compute these using the type 1 SDT model (see Table 1). Using the type 1 SDT model makes the strong assumption that participants use the same decision axis and the same evidence distributions to construct their type 1 and their type 2 judgements. If in an experiment there is, for example, a long latency between making the objective and subjective report, then this might lead to an accumulation of noise (or additional sensory evidence) between the two decision-making stages. This would render the assumption that type 1 and 2 decisions are based upon the same SDT model invalid. Therefore, using this method is only valid when participants have optimal metacognition and, by definition, are using all available type 1 evidence in their metacognitive judgement.

In light of this, we propose using the confidence criteria from the meta-ď fit, *meta*-*c2R1* and *meta*-*c2R2.* Like their *meta*-*ď* counterpart, these criteria represent the confidence criteria an SDT-optimal observer would use to obtain the subject’s empirical responses. Critically, however, these estimates can also be interpreted in another manner: Assuming that the subject is placing confidence criteria rationally, *meta*-*c2R1* and *meta*-*c2R2* can also be interpreted as the confidence criteria over an *alternative* (i.e. enhancement or degradation) model from which the *subject* is constructing their confidence. In other words, the *meta*-*ď* model can be interpreted as the subject’s metacognitive SDT model. On this account, inequality between *meta*-*ď* and *ď* arises either due to additional noise incorporated into the evidence distributions, or more generally due to confidence judgements being constructed from a different decision variable. This could include, for example, an enhanced or degraded signal, or a transformation of the type 1 evidence such as the evidence in favour of the chosen signal class (Maniscalco, Peters, & Lau, 2016). On this interpretation of the *meta*-*ď* model, metacognitive threshold for each response type, *m*-*distanceR1* and *m*-*distanceR2*, are given simply by the difference between the meta-type 1 criterion and the parameters *meta*-*c2R1* and *meta*-*cR2.* This distance is divided by *meta*-*ď* to place it in units comparable across subjects and conditions (see Figure 2).

**Figure 2.**
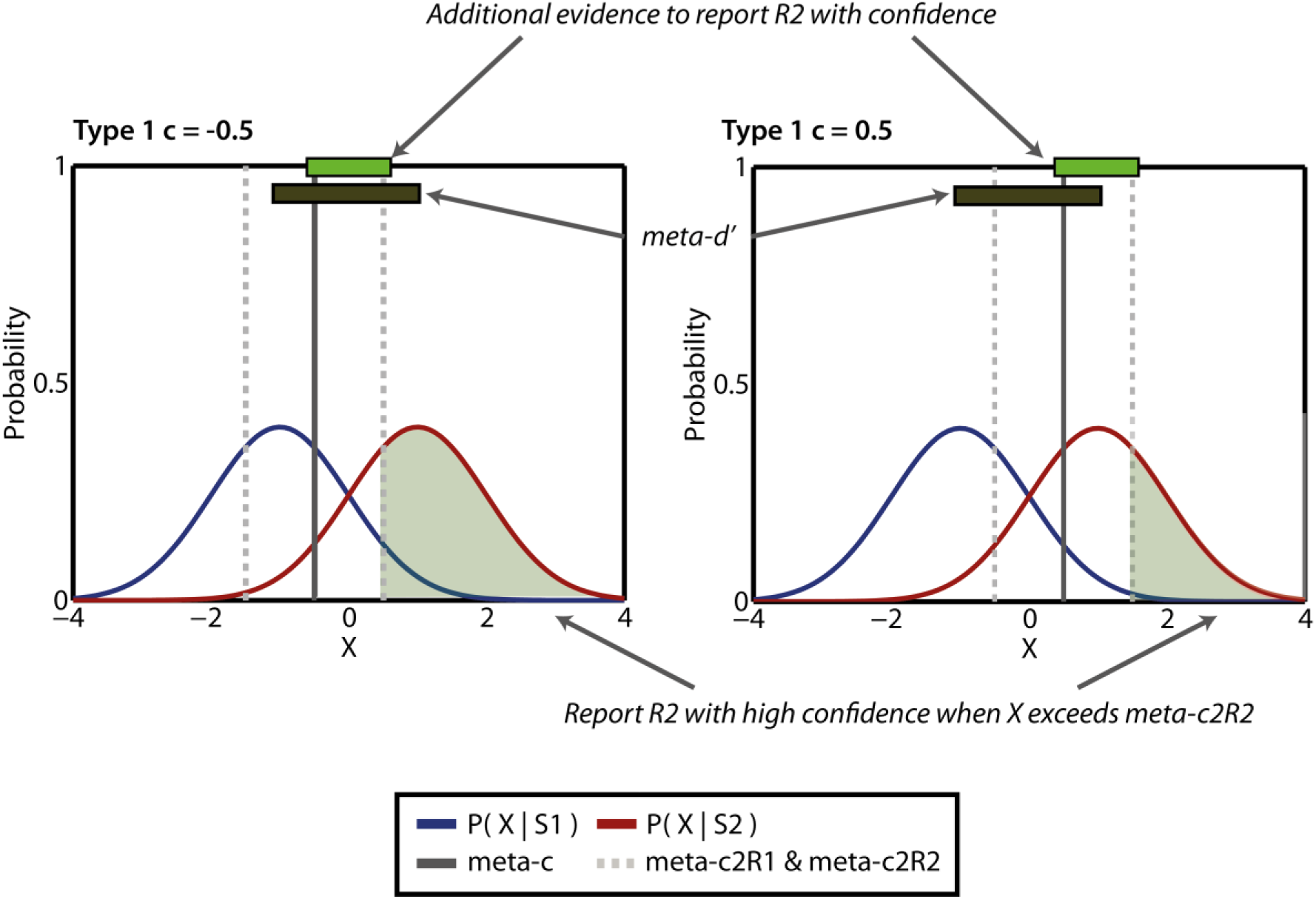
M-distance. The measure m-distance can be computed separately for responses R1 and R2. Metacognitive thresholds meta-cR1 and meta-cR2 are shown in dashed grey lines. Their distance from the meta-c (the type 1 criterion of the metacognitively ideal observer, solid grey line) is depicted by the green bar. M-distance is estimated as a proportion of meta-ď (black bar) so that the measure of metacognitive threshold is independent of metacognitive sensitivity. The shaded green regions depict the values X must take for R2 reports to be given with confidence (independently of the stimulus class). The left and right panels show that, theoretically, m-distance will be invariant under translations of the whole set of type 1 and 2 confidence criteria.

Note that using the *meta*-*ď* model to quantify metacognitive *sensitivity* does not require strong assumptions about the decision variable upon which the metacognitive system operates (Maniscalco & Lau, 2014). This is because the *meta*-*ď* model is not assumed to generate the subject’s decisions. Instead of making this assumption, using the *meta*-*c2* parameters to quantify metacognitive *threshold* does make the alternative assumption that the meta-ď model is involved in generating the subject’s type 2 decisions.

The computations needed to obtain *m*-*distanceR1* and *m*-*distanceR2* are straightforward. The *meta*-*ď* model fixes the type 1 criterion so that *meta*-*c*/*meta*-*ď* = *c*’/*ď*. This results in the *meta*-*ď* model predicting the proportion of *R1* versus *R2* responses exhibited by the empirical data. Therefore, *m*-*distance* is simply given as follows:

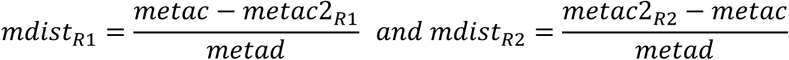

Note that these are standardised by *meta*-*ď* so that *m*-*distance* can be compared across subjects exhibiting different degrees of metacognitive sensitivity.

These equations can also be rewritten as follows:

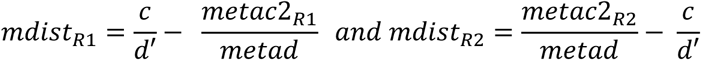

Note that these equations involve subtracting confidence criteria from the (relative) decision threshold. There are two reasons for this. First, the flanking confidence criteria will track changes to the decision threshold. This means that experimental manipulations that induce criterion shifts will also change the flanking confidence criteria without necessarily changing their distance from type 1 criterion. By subtracting the confidence criteria from the type 1 criterion, measures of metacognitive threshold remain invariant to the decision criterion by design, just like their *meta*-*ď* counterpart. The second advantage of this subtraction is that it captures asymmetric changes to the confidence criteria. For example, if an experimental manipulation should induce a preference for one response type over another (e.g. ‘yes’ responses) then *m*-*distance* for ‘yes’ but not ‘no’ responses may change as a function of condition. A single measure of metacognitive strategy that does not consider both types of response (i.e., a ‘response-unconditional’ measure) would not be able to capture any such effect. Of course, when confidence thresholds are assumed to be symmetrically placed about *meta*-*c*, *m*-*distance* reduces to *meta*-*c2R2* – *metac2R1*, normalised by *meta*-*ď*.

## 3. M-distance on empirical data: confidence thresholds are more liberal in participants with above-chance performance

In order to demonstrate the utility of this measure we applied it to published, publically available data by Scott and colleagues (2014). These data comprise responses on an Artificial Grammar Learning (AGL) task, in which participants were first shown a series of letter strings, constructed from an artificial grammar (a set of rules determining allowed sequential transitions between symbols). In the test phase participants were shown some strings whose sequence obeyed this grammar (i.e. were grammatical) and some which did not (i.e. were ungrammatical). Their task was to report (i) whether or not the string was grammatical and (ii) their binary confidence in that judgement. Their metacognitive efficiency indicates the degree to which their grammar learning was implicit versus explicit.

Scott et al. separated participants into one group that had a type 1 ď ≤ 0 in the first 75% of trials (no sensitivity group) and another that had type 1 ď > 0 (sensitivity group). Analyses were conducted on test trials from the remaining 25%. The authors showed that participants in the ‘no sensitivity’ group exhibited metacognitive sensitivity (meta-ď>0) in the absence of type 1 sensitivity. The authors termed this ‘Blind Insight’. Furthermore, the two groups did not significantly differ in their metacognitive efficiency.

We extended this analysis to ask whether these two groups of subjects differed in their metacognitive thresholds, comparing results from *m*-*distance* and prop. confident. We used the same exclusion criteria as Scott et al. (we refer the reader to the original paper), but we further excluded participants with |meta-ď| < 0.1 in the selection trials (meta-ď_selection_). This was to minimise the frequency with which we divided by very small values when computing m-distance. Type 1 ď in the selection trials did not significantly differ between participants with meta-ď_selection_ < 0.1 and meta-ď_selection_ ≥ 0.1, t(433) = 1.59, *p* = .110, so this additional exclusion criterion did not bias the sample. We used bootstrapped independent t-tests (10,000 samples) to determine whether the differences between groups were significant. Because they are bounded variables, both the proportion of ‘confident’ responses and m-distance were log-transformed.

As depicted in Figure 3, both measures unsurprisingly revealed the same trend: a greater propensity to report with confidence when participants had type 1 sensitivity. However, while this was not significant when using (log) proportion confident, *t*(70.63) = 1.87, *p* = .061, the difference was significant under log(m-dist), t(410) = 2.21, *p* = .027. Therefore, participants who exhibited type 1 sensitivity to the task (i.e. were able to discriminate grammatical from ungrammatical strings) had more liberal confidence thresholds.

**Figure 3.**
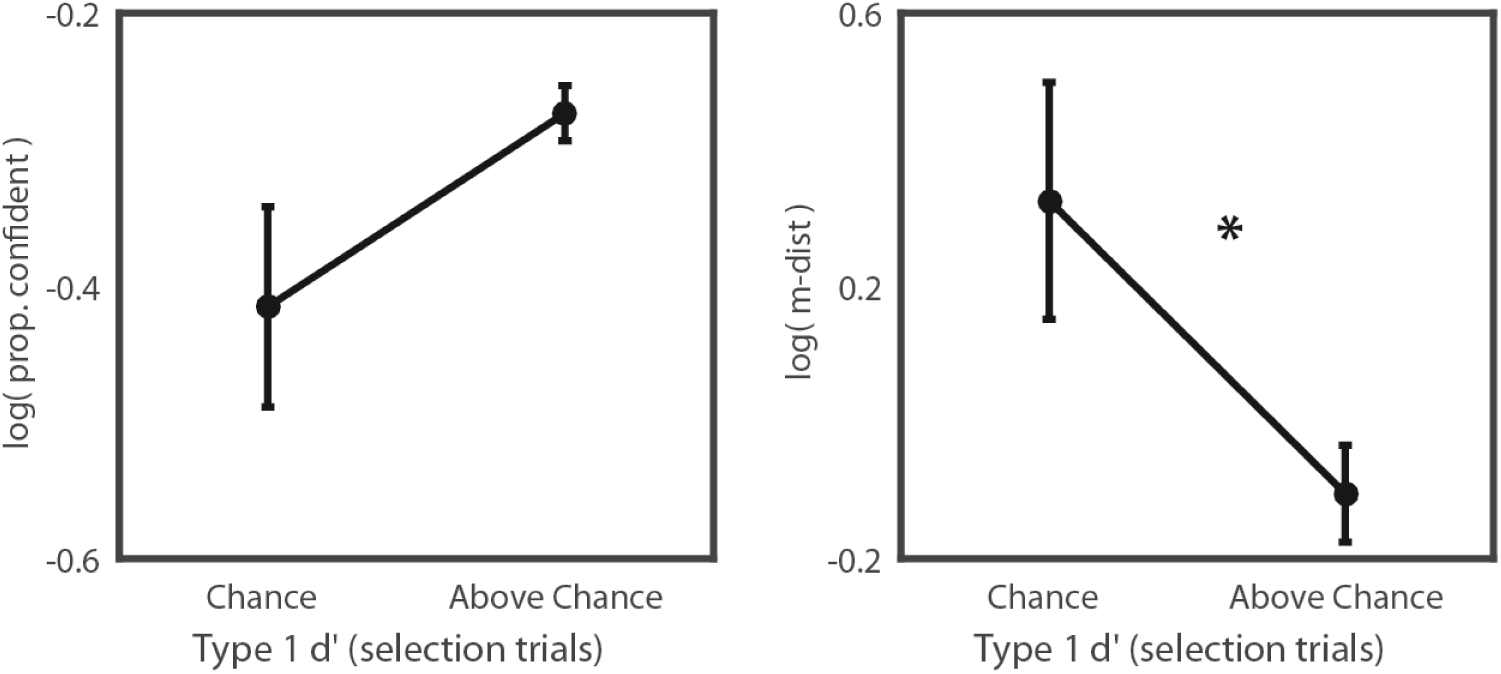
Confidence and m-distance (data from Scott et al, 2014.) for test-trial data (last 25%) The left panel shows that the proportion of confident responses is higher in participants who showed type 1 sensitivity in selection trials than in those who did not. This effect was non-significant. The right panel shows that m-dist was reduced, i.e. confidence criteria were closer together in participants who showed type 1 sensitivity in selection trials. Therefore, participants with type 1 sensitivity had a greater propensity to report with confidence. This effect was statistically significant. ^⋆^ p < .05

In summary, *m*-*distance* was able to detect a statistically significant effect that proportion confident was not. This is important, because while using confidence ratings would lead one to conclude that participants gave similar confidence reports regardless of type 1 performance, the metric m-distance reveals that in fact these groups behaved differently in their confidence judgements. We note that this analysis does not undermine the central claim in Scott et al., namely the ‘no sensitivity’ group showed significantly non-zero metacognitive sensitivity.

## 4. Simulations

### 4.1 Methods

Having derived a principled measure of metacognitive thresholds, *m*-*distance*, and tested it on an illustrated empirical dataset, we further examined its empirical behaviour by simulating responses to 100,000 trials under a range of conditions, using a generative model based on SDT assumptions. We also compared its behaviour with that of the proportion of trial responses made with confidence (prop. confident, Fig. 1A). Unless stated otherwise, trials were simulated from an equal-variance SDT model with *ď* = 2, *c* = 0 (i.e. an unbiased observer), meta-*ď* = 2 (i.e. optimal metacognitive performance) and flanking confidence criteria *c2*_R1_ and *c2*_R2_ which extended 0.5 units out from *c.* Under these parameters m-distance for each response type is 0.25. The response-conditional meta-*ď* model was estimated using Maniscalco and Lau’s method, which relies on maximum likelihood estimation (Maniscalco & Lau, 2012).

### 4.2 Results

#### 4.2.1 M-distance as a function of type 1 and 2 thresholds

First we verified that *m*-*distance* is indeed robust to changes in type 1 criterion. Criterion *c* was varied between −1.5 and 1.5 in steps of 0.1. The distance between *c* and C2_R1_ and C2_R1_ was fixed at the default of 0.5, but the numerical values they took were *c* - 0.5 and *c* + 0.5 respectively. All other parameters were set to their default values (table 2).

**Table 2.**
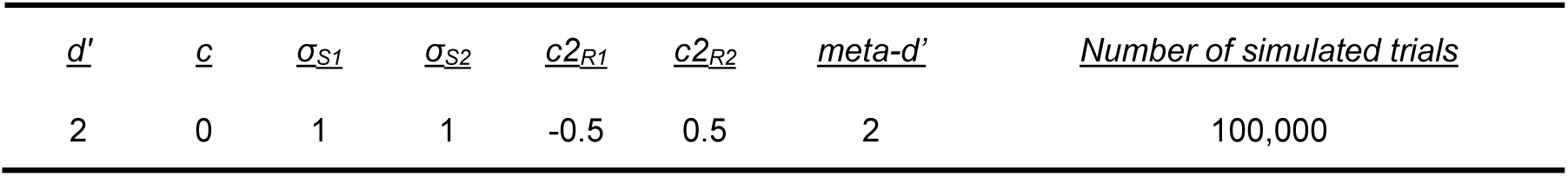
Default parameter values

| $d'$ | $c$ | $\sigma_{S1}$ | $\sigma_{S2}$ | $c2_{R1}$ | $c2_{R2}$ | $meta-d'$ | *Number of simulated trials* |
| --- | --- | --- | --- | --- | --- | --- | --- |
| 2 | 0 | 1 | 1 | -0.5 | 0.5 | 2 | 100,000 |

As shown in Figure 4A, while prop. confident was strongly biased by changes in type 1 *c*, m-dist stayed constant with respect to criterion, though became somewhat unstable at extreme values.

**Figure 4.**
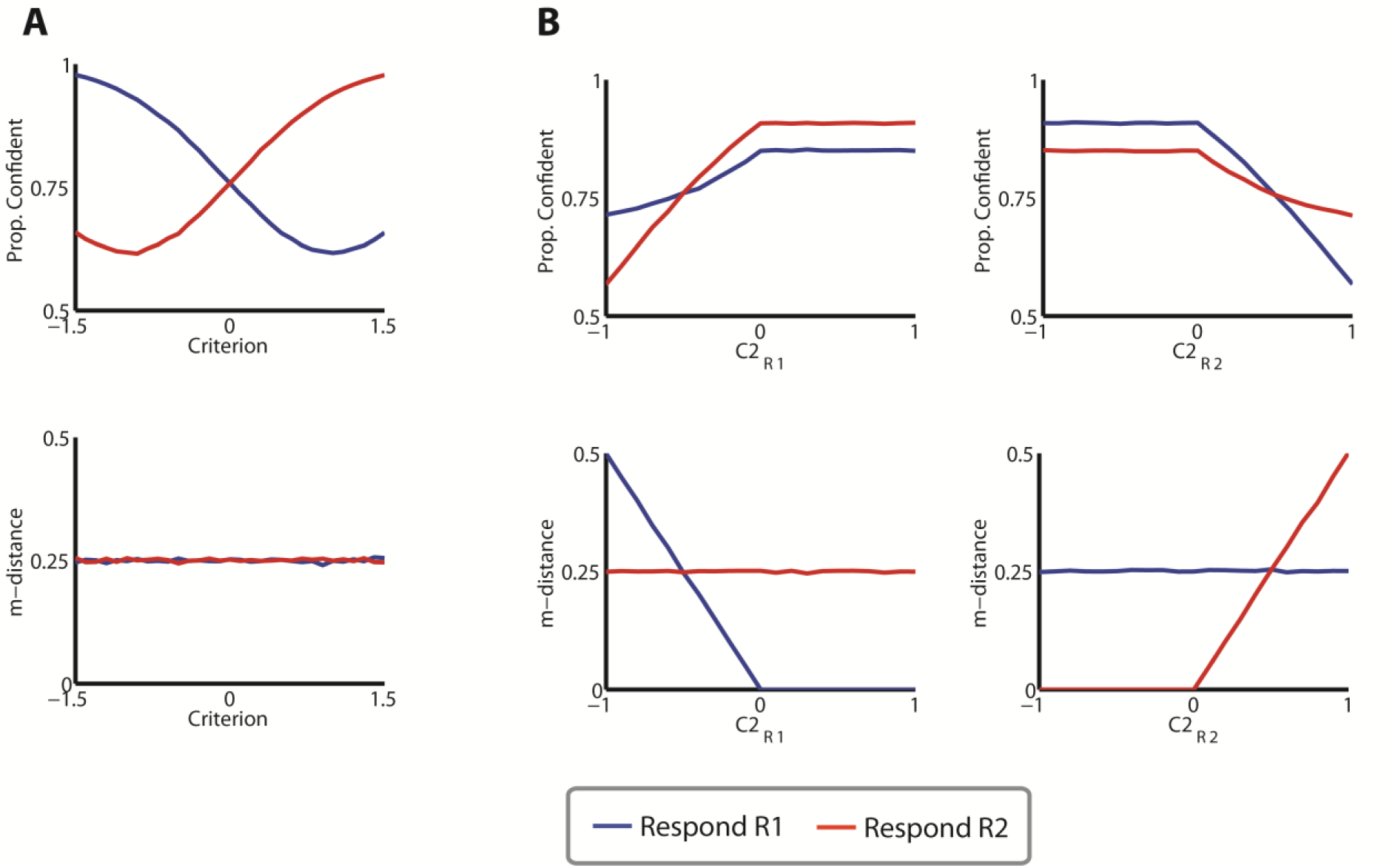
Proportion of confident responses and m-distance for R1 (blue) and R2 (red) responses, as a function of changing SDT parameters. **A.** M-distance (bottom) but not prop. confident (top) is stable under changing type 1 criterion. **B.** Prop. confidence (top) and m-distance (bottom) as a function of C2_R1_ (left) and C2_R2_ (right)

Next, we verified that changes to one of the confidence criteria would result in changes to the corresponding *m*-*distance* value. Type 1 criterion *c* was fixed at zero. First, we fixed *c2*_R2_ at 0.5 and varied *c2*_R1_ between −1 and 1 (figure 4B). For R1 responses, both prop. confident and *m*-*distance* were invariant to this parameter, as expected. Also as expected, more extreme values of *c2*_R1_ were associated with more extreme values of *m*-*distance* for R1 responses (i.e. more conservative confidence criteria) and lower confidence. When *c2*_R1_ surpassed the type 1 threshold, all trials were classed as ‘confident’ and so m-distance was maximally liberal (i.e. was equal to zero). We repeated this, fixing *c2*_R1_ at −0.5 and varying *c2*_R2_ between −1 and 1 units away from criterion (figure 4B, right). The analogous result was found.

It is important to note that some experimental paradigms are thought to violate the equal variance SDT model (EVSDT). For example, recognition memory yes/no tasks, the memory trace for a previously seen stimulus ought to be more precise than that for a novel stimulus. Accordingly, these tasks are thought to be best fit by unequal variance SDT (UVSDT) (Mickes, L, Wixted, J.T., Wais, 2007). M-distance values will be confounded if it is estimated assuming EVSDT when the equal variance assumption has been violated. This is because different evidence distributions with fixed criteria will generate different values of m-dist.

In summary, m-distance is preferable to prop. confident in experimental designs where type 1 criterion is being modulated as long as EVSDT is not incorrectly assumed.

#### 4.2.3 M-distance as a function of type 1 sensitivity

It is clear from empirical work that confidence increases with task performance (Grimaldi et al., 2015), and for this reason it is often advised to equate performance across conditions if one is probing confidence. Our next simulation illustrates how *m*-*distance* changes with type 1 *ď*. We varied type 1 *ď* between 0.5 and 3 in steps of 0.1. All other parameters kept their default values (table 2, Figure 5A and 5B). As shown in Figure 5C, confidence (as measured by prop. confidence) increases with increasing type 1 *ď*. Similarly, and as shown in Figure 5D, *m*-*distance* decreased (i.e. there was a greater propensity to report ‘confident’) as type 1 *ď* increased. In other words, given fixed criteria, better objective sensitivity led to smaller values of *m*-*distance* and increased tendency to report ‘confident’. As can be seen from the type 1 SDT models from which these trials were simulated (see Fig 5A and 5B), despite the simulated subject requiring the same amount of absolute evidence X to report with confidence, they are more likely to exceed that amount when *ď* is high.

**Figure 5.**
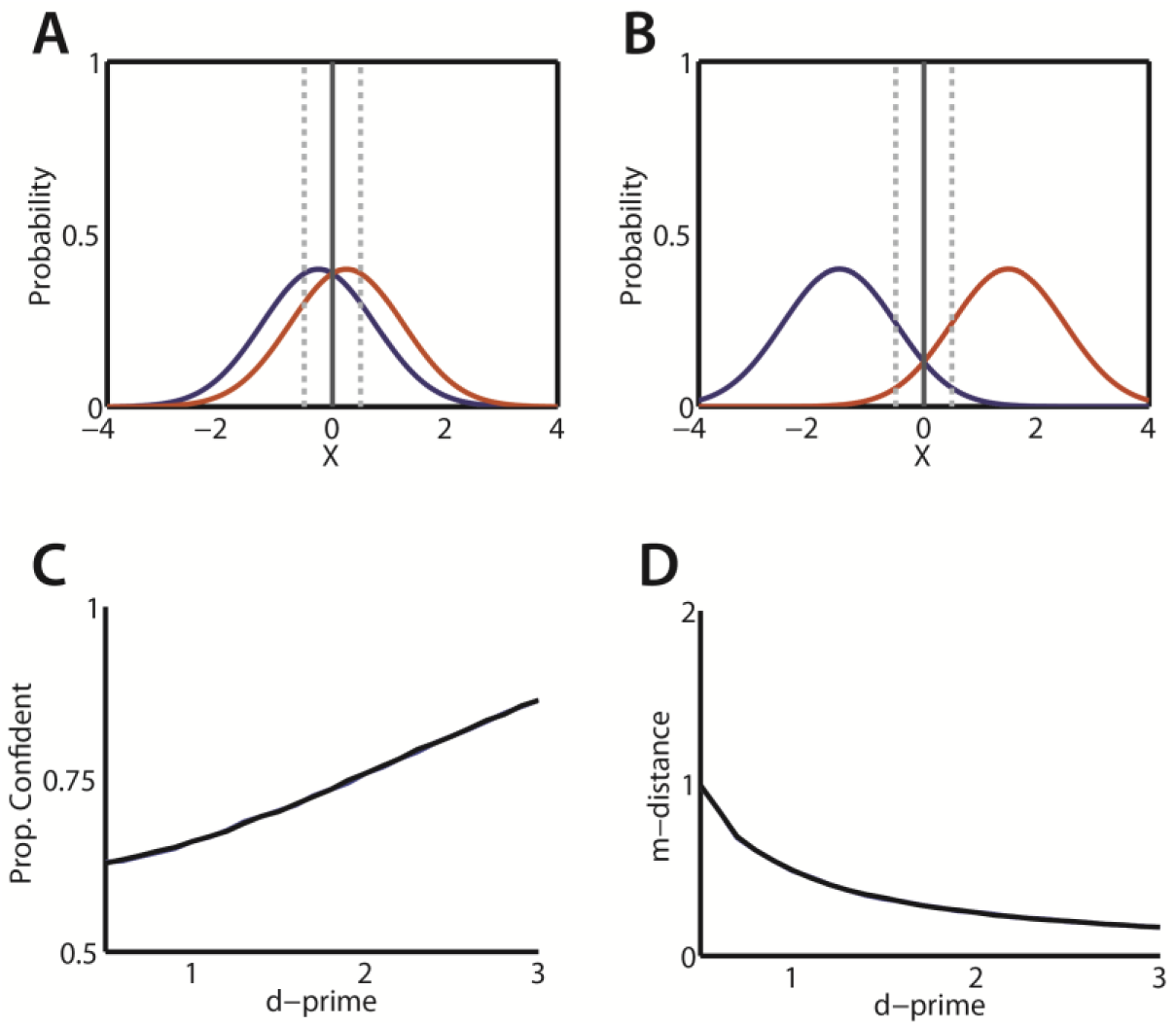
**A, B.** The (type 1) SDT models from which trials were simulated when **A.** ď = 0.5 and when B. ď = 3. **C.** Prop. confident as a function of type 1 ď. **D.** M-distance as a function of type 1 ď.

#### 4.2.4 Sensitivity of *m*-*distance* to trial count

To determine the sensitivity of m-distance to trial count, we replicated the simulations in which the type 1 decision threshold was varied in the case where *meta*-*ď* = *ď*, but now also with varying numbers of simulated trials. We simulated responses from 10 observers completing between 100 and 1000 trials (in intervals of 100). We varied *c* such that it took values between −2 and 2 in each case. We set *ď* = 2 and placed confidence thresholds symmetrically about the decision criterion, extending out by 0.5 evidence units. If the measure is stable then for both response classes, the stimulated value of *m*-*distance* should be equal to 0.5. The stability will depend upon whether enough trials of each class (e.g. confident false alarms) are present: when *c* takes extreme values relative to *ď* it is more likely that there will be cells with a low trial count, leading to unstable estimates when computing the Z-transform of the hit rate and false alarm rate. Accordingly we followed the common procedure of adding 0.5 to each cell count (confident hits, guess hits etc.) so that SDT quantities can still be computed on otherwise empty cells.

Figure 6 depicts the values of *m*-*distance* obtained for each simulated subject and each number of trials. Some simulated subjects continued to exhibit substantial deviations from the expected value of 0.5 until the number of trials collected reached 400 per condition. However under extreme criteria, when | *c* | > 0.75*ď*, deviations from the expected value persisted even with 1000 trials.

**Figure 6.**
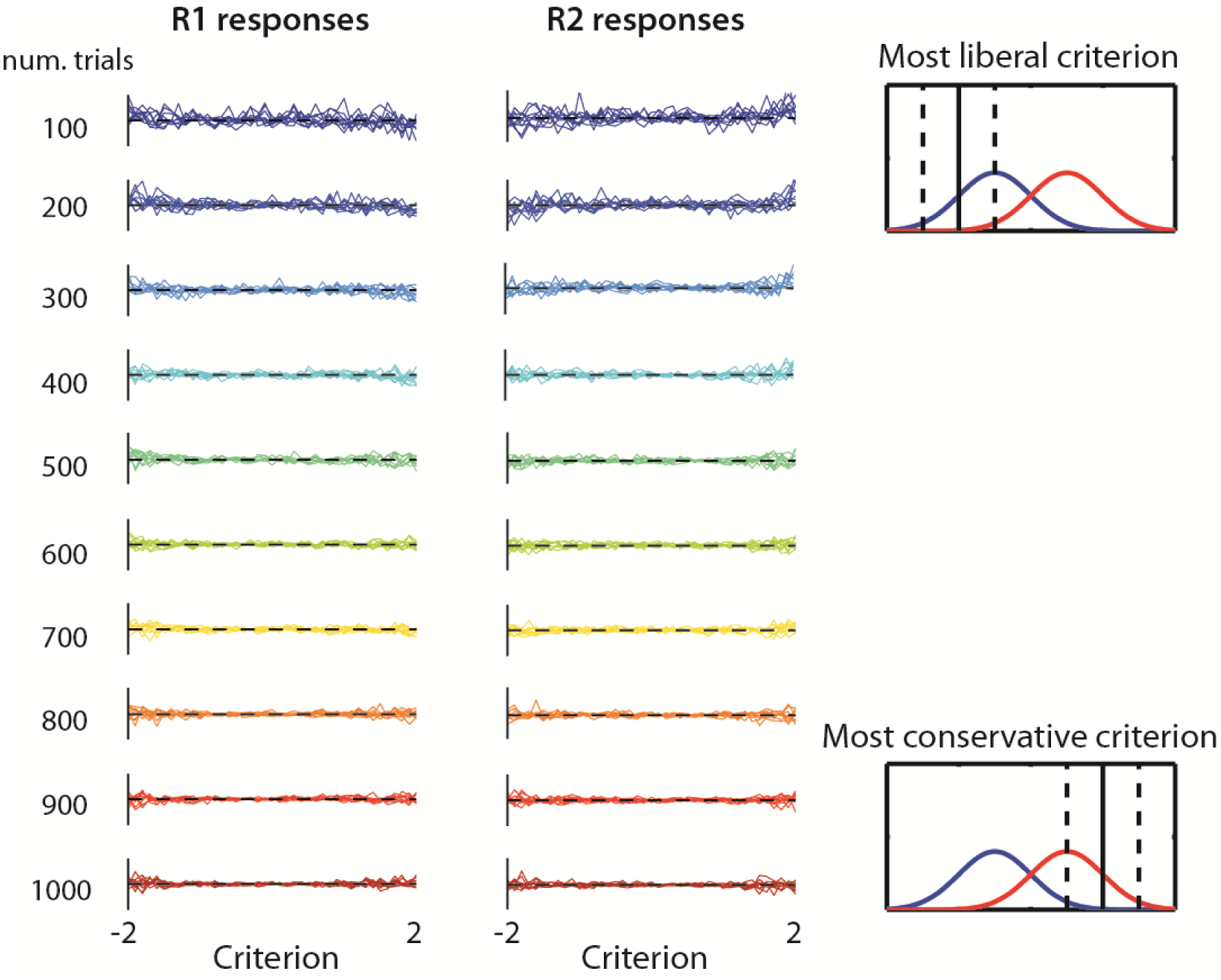
Simulated values of m-distance as a function of decision criterion, response class (column) and trial count (row). Each panel depicts the estimated m-distance value over the range of decision criteria tested for 10 simulated subjects. The y-axis goes from 0 to 1. Colours represent the trial count (where dark blue is the lowest and dark red is the highest). The right-most panels illustrate SDT models from which trials were simulated. Specifically, the top-right panel illustrates the SDT model with the most liberal criterion (c = −2, c’ = −1) and the bottom-right panel illustrates the model with the most conservative criterion (c = 2, c’ = −1).

#### 4.2.5 Alternative SDT models: Interpreting meta-*ď*/*ď* as the signal change between objective and subjective reports

Why might metacognitive sensitivity deviate from SDT-optimality? Signal degradation models assume that after the objective judgement has been made the evidence degrades because either time has elapsed or because of additional internal noise. Such a model can account for conditions under which confidence is less sensitive to some manipulation than the objective judgement (e.g. when above chance performance but zero confidence is observed, Peirce and Jastrow 1884). In these cases, despite objective performance remaining constant across conditions, confidence decreases, in turn leading to reduced metacognitive sensitivity.

In some cases, empirical data reveal values of *meta*-*ď* that exceed *ď* (Rahnev & Denison, 2016). Metacognition that exceeds that of the SDT-optimal observer can be modelled under a signal enhancement model, where evidence increases (or noise decreases) between the systems instantiating the type 1 and 2 judgements. For example, in an artificial grammar learning task, above-chance metacognition can be observed in the absence of any type 1 sensitivity (Scott et al., 2014). This could be modelled via additional (e.g. top-down) sources of evidence that only target the subjective decision. In SDT terms, this would correspond to a decision axis that better distinguishes the likelihood functions.

Figure 7 illustrates some alternative SDT models. The aim of these is to account for these dissociations between type 1 and type 2 decisions. These alternative SDT models are not mutually exclusive proposals for how confidence is constructed. Rather, they explain the different relationships between confidence and accuracy that may arise under different task conditions. That is, a signal enhancement model may capture confidence judgements on one task (e.g. where subjects receive feedback on their choice accuracy prior to reporting confidence), while performance on another task might be better captured by a signal degradation model (e.g. where there is a long latency between reporting choice and confidence).

**Figure 7.**
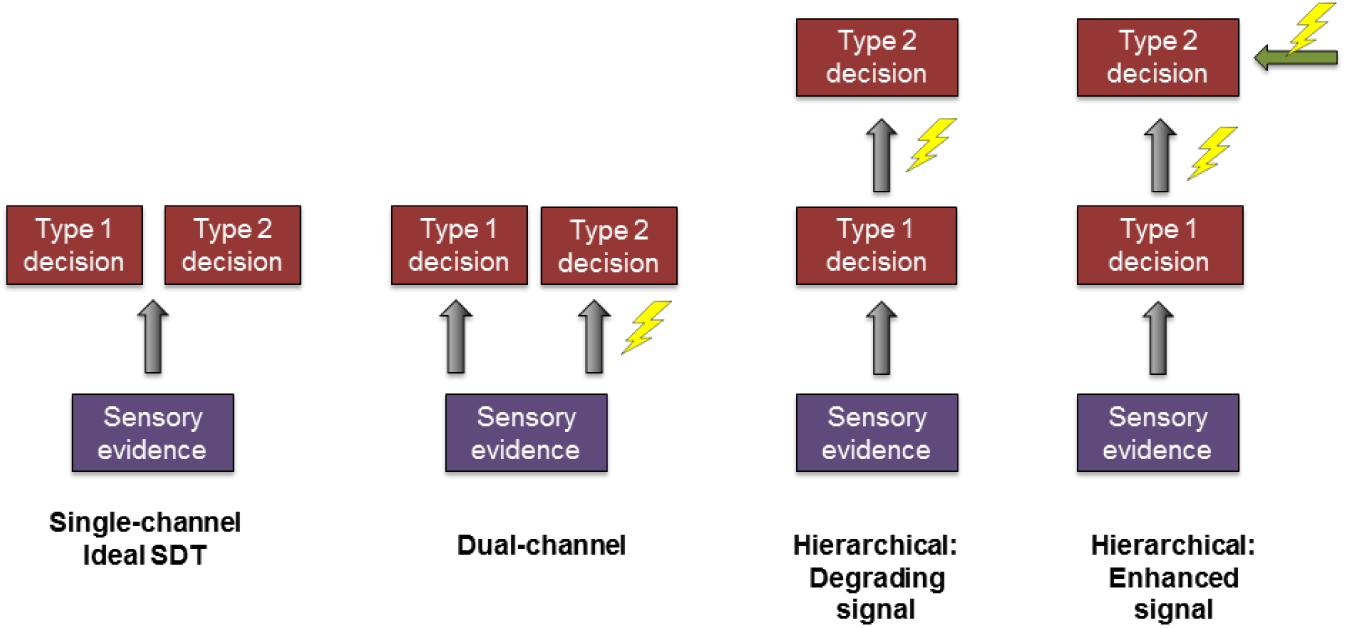
Alternative models of metacognition. From left: single-channel, which assumes that the perceptual and metacognitive systems receive the same input; dual-channel, which assumes that each system receives independent (or partially independent) representations of the same source; and hierarchical models, which assume that the metacognitive system receives the outputs of the perceptual system. Lightening bolts indicate the sources of noise (grey areas) which can be targeted such that metacognition will change. [adapted from Maniscalco & Lau, (2016)]

#### 4.2.6 Metacognitive parameters under hierarchical models

We implemented a hierarchical model as follows. For some evidence *x_k_* received on trial *k* (from which the objective decision is made), the evidence made available for the confidence report *y_k_* is given by (Barrett et al., 2013):

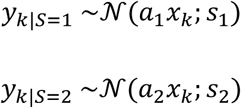

Here, *s*_1_ and *s*_2_ are independent free parameters (bounded by zero) that determine the degree of additional noise applied to the evidence available for confidence. Additionally, *a*_1_ and *a*_2_ are independent free parameters bounded by zero which scale the evidence. When *a*_1_,*a*_2_ <1 we have a signal degradation model, in which the absolute evidence available for confidence, *y_k_*, is reduced. Here, the evidence regresses towards the intersection of the two likelihood functions. This might represent, for example, signal decay over time (e.g. forgetting). When *a*_1_,*a*_2_ > 1 the evidence available for confidence is amplified and pushed towards the extremities. This could represent continued exposure to the stimulus after the type 1 report.

Note that when *a*_1_= *a*_2_= 1 and *s*_1_= *s*_2_ = 0 there is no signal change and *meta*-*ď* = *ď*.

To determine the behaviour of *m*-*distance* when *meta*-*ď* ≠ *ď* we set *a*_1_ = *a*_2_ and varied them between 0.5 (shrinking evidence) and 1.5 (amplifying evidence) in steps of 0.1. We set *s*_1_ = *s*_2_ and varied them between 0 and 2 in steps of 0.1. All other parameters were set to our defaults (table 2). In simulations implementing alternative SDT models, the evidence upon which type 1 and 2 decisions are based can fall on opposite sides of the type 1 decision threshold. This would reflect ‘change of mind’. Where this situation arose, confidence was set to zero.

As shown in Figure 8A (top row), *m*-*ratio* (*meta*-*ď*/*ď*) was primarily influenced by *s*, such that more noise was associated with greater deviance from SDT-optimality. This interacted with *a*, such that boosting the evidence (increasing *a*) preserved metacognitive sensitivity under greater levels of noise.

**Figure 8.**
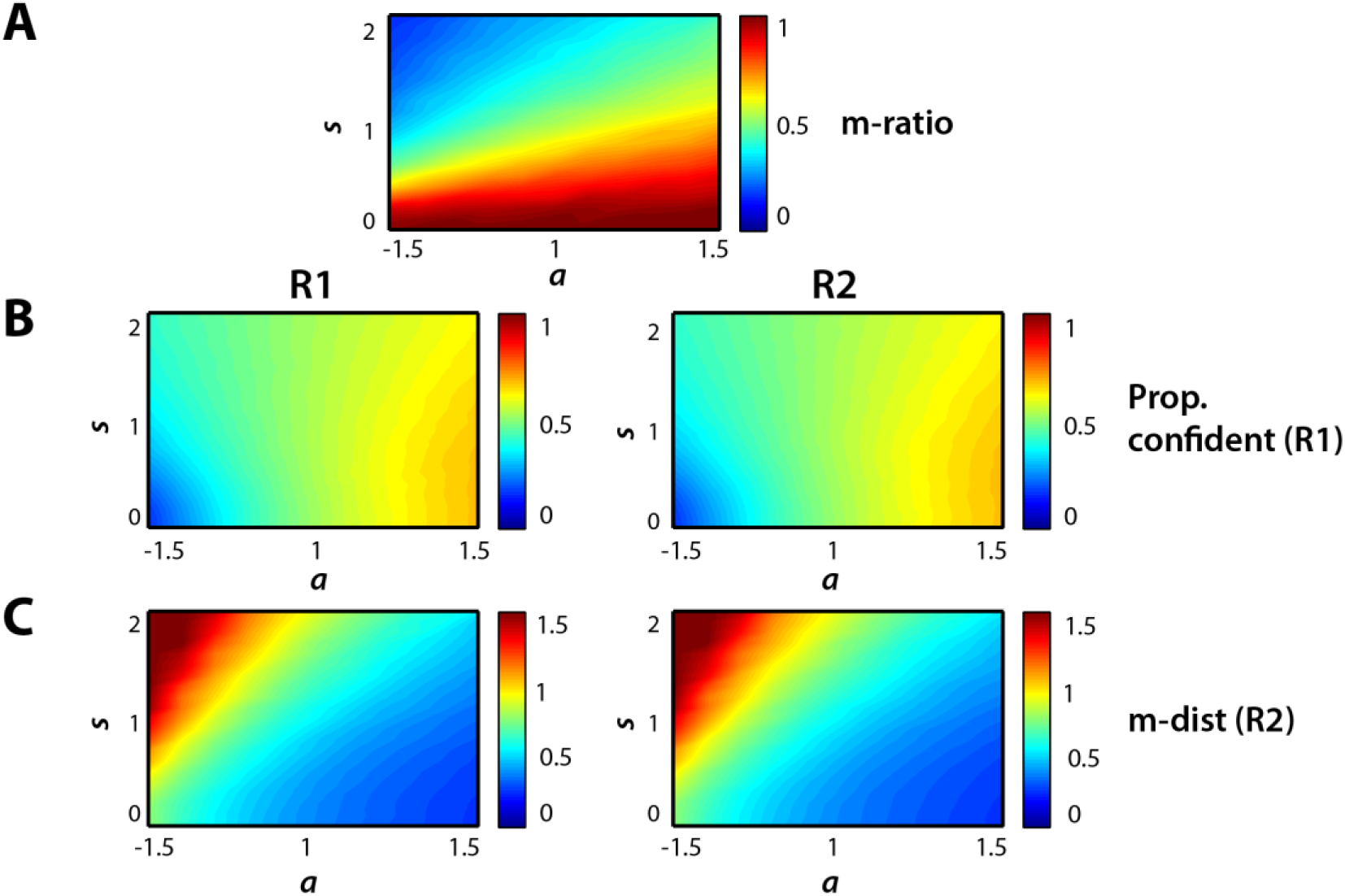
Metacognitive sensitivity and metacognitive thresholds under a hierarchical SDT model. The panels depict **A.** m-ratio, **B.** the proportion of confident responses in cases R1 (left) and R2 (right) and **C.** m-distance for R1 (left) and R2 (right) responses. Values are given as a function of the signal shrinkage a (x-axis) and additional variance s (y-axis).

Prop. confident (Fig 8B) was not particularly affected by these changes to the signal available for metacognition unless the parameter *a* was extreme. In other words, confidence was not very sensitive to signal amplification or shrinkage. This is problematic, because metacognitive thresholds (here assessed with prop.confident) should reflect the amount of additional evidence required to report a decision with confidence.

On the other hand, *M*-*distance* (Fig 8C) exhibited a similar pattern to m-ratio, but the parameter *a* affected it to a greater degree. What this means is that changes to the absolute value of the evidence had a greater effect on confidence than on metacognitive sensitivity.

This simulation shows that hierarchical models can reproduce empirical findings of suboptimal metacognition. However, this model could not reproduce values of *m*-*ratio* > 1, i.e. metacognitive efficiency that exceeds standard SDT predictions (within a reasonable range of parameter choices explored). Therefore, the particular hierarchical model used here cannot be readily applied to tasks which result in *m*-*ratio* > 1.

#### 4.2.7 Behaviour of meta-ď and m-distance under dual-channel models

The simulations above assumed a hierarchical model of metacognition. By contrast, dual-channel models (e.g. Del Cul et al. 2009; Jacoby 1991) assume that type 1 and 2 decisions are subject to independent sources of internal noise. In SDT terms, this is equivalent to positing independent decision axes for type 1 and type 2 judgements, so that the internal noise affecting type 1 judgements is independent from the internal noise affecting type 2 judgements. This is modelled by sampling evidence for the type 1 and type 2 judgements independently. To model signal enhancement or degradation in the dual-channel case, evidence for the type 2 judgement is drawn from likelihood functions with a higher or lower variance than those generating the type 1 judgement.

To test whether the measures *meta*-*ď* and *m*-*distance* remain robust to type 1 criterion changes under dual-channel models, we simulated 100,000 trials from an observer with *ď* = 2. The dual-channel model was implemented by assuming two separate EV SDT models for the type 1 and 2 decisions. The evidence for the type 1 decision had a mean of −1 and a variance of 1 for stimulus S1, and mean 1 and variance 1 for stimulus S2. Reflecting the same sensory signal feeding into both type 1 and 2 decisions, the means of the evidence distributions were unchanged for the type 2 decision. The variances of the S1 and S2 distributions for the type 2 decisions, σ_1_ and σ_2_, were varied together between 0.2 (reflecting a more precise signal for confidence, i.e. signal enhancement) to 2.2 (reflecting a noisier signal for confidence, i.e. signal degradation) in steps of 0.2.

We also varied type 1 criterion between −1 and 1 (in intervals of 0.1). If the *meta*-*ď* and *m*-*distance* behave similarly under this dual-channel model and hierarchical models, the measures should remain largely constant with respect to type 1 *c*. The two confidence criteria were placed 0.5 evidence (X) units away from type 1 *c*.

Figure 9A depicts the behaviour of *m*-*ratio* as a function of type 1 *c* and the variances of the type 2 (metacognitive) distributions. These results show that *meta*-*ď* is not entirely invariant to changes to type 1 *c* under the dual-channel model implemented here. Specifically, *M*-*ratio* decreased as the decision threshold became more biased. Effects of type 1 *c* were also seen in a more exaggerated form for both prop. confident (Fig 9B) and *m*-*distance* (Fig 9C).

**Figure 9.**
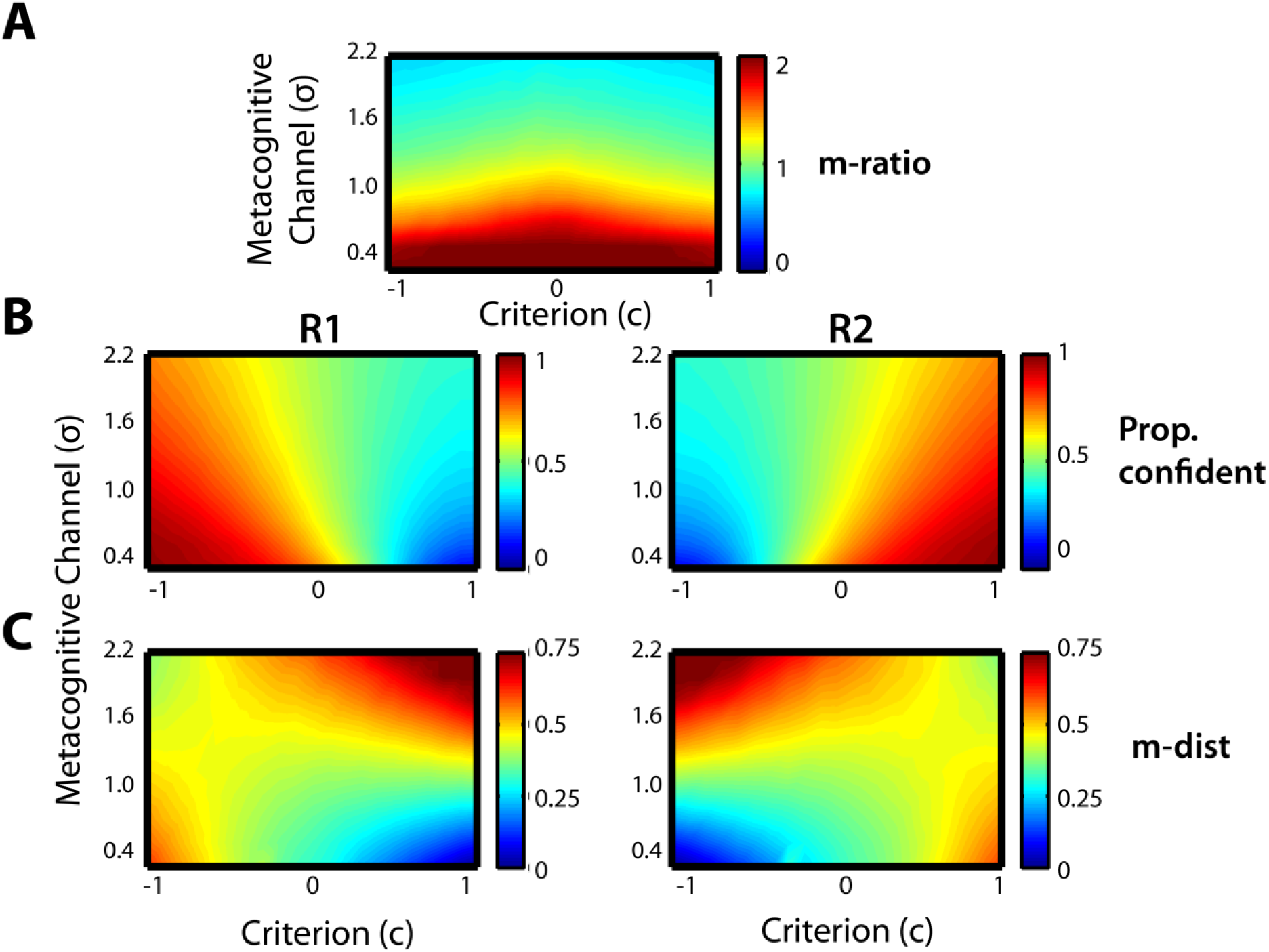
Behaviour of m-distance and m-ratio under a dual-channel model. **A.** Behaviour of m-ratio under a dual-channel model. M-ratio is not invariant to criterion under this model. The measure decreases as criterion becomes increasingly biased (i.e. deviates from zero). It also tracks σ, such that under signal enhancement (σ < 1) metacognitive sensitivity surpasses that of the SDT-optimal observer and under substantial signal degradation (σ > ~1.6) metacognitive sensitivity is less than that of the SDT-optimal observer. **B.** Prop. confident for R1 and R2 responses. Increasing noise σ somewhat reduces the effect of type 1 c on confidence, and overall, greater noise is associated with lower confidence as would be expected. **C.** M-distance for R1 and R2 responses. M-distance is somewhat robust to type 1 c under these models, except under extreme criteria or degradation. Furthermore, m-distance strongly increases with signal degradation.

Similarly to what was found for hierarchical models (Figure 8), greater degradation of the confidence signal (increasing σ) caused *m*-*ratio* to decrease. However, hierarchical and dual-channel models diverged in two key ways. First, while the hierarchical model could not reproduce values of *m*-*ratio* > 1 (without rather extreme parameter choices), this dual-channel model could. Second, while the hierarchical model did reproduce values of *m*-*ratio* < 1, the dual-channel model required a rather large amount of noise in the metacognitive channel for *m*-*ratio* to fall below 1.

As shown in Figure 9C, a more precise signal for confidence (σ < 1) led to lower values of *m*-*distance* (greater propensity to report ‘confident’) whereas a noisier signal for confidence led to higher values of *m-distance.* When σ was either very large or very small, *m*-*distance* lost its invariance to type 1 criterion when that criterion was extreme. In particular, an extreme criterion can lead to hit rate and false alarm rate approaching 0 or 1. At these extreme values, their z-transforms become unstable. Confidence (Figure 9B) exhibited a similar pattern, but was less robust to changes in type 1 *c* and less sensitive to changes in the confidence signal.

The above simulations do not allow full determination of the circumstances under which *meta*-*ď* should be used as an estimate of metacognitive sensitivity. Ideally, behaviour of the measure should be tested on simulations of plausible models for each new application. However, the hierarchical and dual models that we have considered are reasonable in a considerable number of realistic scenarios. We conclude from these simulations that, for these models, *meta*-*ď* will be robust if the type 1 criterion doesn’t vary too much between conditions, and that the new *m*-*distance* confidence measure is robust to changes in both type 1 criterion and *meta*-*ď*. Furthermore, we show that *m*-*distance* is preferable to proportion confident when subjects exhibit a decision bias and/or *meta*-*ď* differs from *ď*.

## 5. Discussion

Investigations of subjective confidence, and of metacognition more generally, require empirically robust and theoretically principled measures of metacognitive threshold. Using the signal-detection-theory framework of *meta*-*ď*, we have defined and quantitatively characterised a novel measure of metacognitive threshold – *m*-*distance* - that is theoretically robust to changes in both type 1 criterion and metacognitive sensitivity. *M*-*distance* reflects the amount of internal evidence required to report a forced choice (e.g. “left” versus “right”) with confidence, relative to the amount of evidence needed to make that choice in the first instance.

Lower values of *m*-*distance* indicate more liberal confidence criteria, because the subject only needs a small amount of additional evidence to report their choice with confidence. The reverse applies for conservative criteria. Given minimal assumptions about how metacognitive decisions are made (discussed in the next paragraphs) and with a sufficient number of trials per condition (approximately 300), this measure is empirically accurate and indexes metacognitive thresholds in a meaningful manner on stimulus classification tasks.

It is of note that *m*-*distance* is not a measure of confidence *bias*, i.e. towards being over- or under-confident. Type 1 *c* has a clear reference point at which the two response types require equal amounts of evidence: the intersection of the two likelihood functions. Accordingly, we can talk about type 1 decisions being biased towards R1 or R2. In contrast, metacognitive thresholds *meta*-*c2R1* and *meta*-*c2R2* do not have a reference point above which the subject is more confident than is warranted by their type 1 performance. We leave the question of how best to identify such a reference point open for future research.

Further simulations revealed that *m*-*distance* behaves as expected on alternative SDT models, including hierarchical models where the signal degrades or is amplified between the type 1 and 2 decision, as well as on dual-channel models where evidence is sampled separately for type 1 and 2 decisions. This means that strict assumptions about how metacognition is achieved at the computational level do not need to be made for *m*-*distance* to be a meaningful and valid measure of metacognitive threshold. Put more simply, *m*-*distance* is a useful measure of propensity towards reporting with low (or high) confidence across a range of possibilities for how metacognitive decisions might be implemented in the brain.

Our simulations replicated previous work that, in hierarchical models, suboptimal metacognition (i.e. *meta*-*ď* < *ď*) can arise either because additional internal noise is integrated into the signal used by the metacognitive system, or because the signal magnitude decreases (Barrett et al., 2013). Since that work, alternative dual-channel models have been highlighted as possible implementations of metacognition (e.g. Timmermans, Schilbach, Pasquali, & Cleeremans, 2012). In the present paper we additionally tested the behaviour of *m*-*ratio*, prop. confident and *m*-*distance* on a dual-channel model, in which fully independent sources of internal noise are integrated into the type 1 and 2 judgements. These simulations revealed that *meta*-*ď* loses its invariance to type 1 criterion when confidence is constructed this way: *m*-*ratio* will decrease as type 1 criterion becomes increasingly biased. Thus, *m*-*distance*, because it is a function of *meta*-*ď*, will be biased by criterion shifts. However, type 1 c has a smaller effect on *m*-*distance* than on prop. confident, rendering m-distance a better choice for estimating confidence in tasks amenable to SDT.

Crucially, *m*-*distance* was more sensitive to changes in the signal to noise ratio available to the metacognitive sensitive than prop. confident was. This indicates that *m*-*distance* is a better measure, because confidence should reflect the amount of additional evidence required for ‘confident’ report: changes to the available evidence should be reflected in the measure of confidence used.

Curiously, while hierarchical models could not recreate the ‘super-optimal’ metacognition sometimes reported in the literature (Rahnev & Denison, 2016), dual-channel models could: using the parameters here, *meta*-*ď* exceeded *ď* by a factor of up to (and even over) two. However, increasing the variance of the distributions upon which confidence judgements were based (the σ parameter) did not reduce *m*-*ratio* as dramatically. These simulations suggest that a fruitful approach to modelling construction of confidence in an SDT framework may be to construct a hybrid of these two families of models.

By using *m*-*distance* as an alternative computation to mean confidence, research into subjective confidence and metacognition will be better equipped to interpret confidence judgements in terms of changes to putative metacognitive thresholds. For example, previous work has found that mean confidence increases for perceptual decisions that are congruent with prior expectations (Sherman, Seth, and Kanai 2016). However, the experimental manipulation in this study also biased decision threshold. Therefore, the reported changes in mean confidence may have arisen indirectly as a result of this type 1 criterion shift. Similarly, while confidence usually decreases with increasing sensory noise (Baranski & Petrusic, 1998; Spence et al., 2015), previous work has shown that physiologically arousing cues can render confidence (as measured by *meta*-*c2)* blind to sensory noise (Allen et al., 2016). The use of *m*-*distance* would also have had utility there, by controlling for effects of arousing cues on metacognitive sensitivity *meta*-*ď*.

***

## Summary

In summary, we have introduced a new signal-detection-theoretic measure *m*-*distance.* This is a theoretically principled and empirically robust measure of metacognitive threshold, which is normalised with respect to the agent’s metacognitive sensitivity. In this way, the measure can be compared across subjects and across experimental conditions when metacognitive sensitivity *meta*-*ď* is changing. This is in contrast to using raw prop. confident as a measure, which is less sensitive to signal degradation and enhancement. Using simulations we showed that the confidence criteria arising from the *meta*-*ď* model fit can be used to characterise metacognitive thresholds – which define *m*-*distance* - in a manner that is approximately independent of metacognitive sensitivity and decision thresholds. Furthermore, we have shown that the measure behaves in predictable ways when the signal used for the confidence judgement is either degraded or enhanced. Given a sufficient number of trials (approximately 300 per condition), empirical research into subjective decision-making can therefore use *m*-*distance* to better understand how subjective reports of confidence are constructed in the brain and expressed in behaviour.

## Acknowledgements

The authors are grateful to the Dr. Mortimer and Theresa Sackler Foundation, which supports the Sackler Centre for Consciousness Science and funds MTS. ABB is funded by EPSRC grant EP/L005131/1. AKS also acknowledges support from the Canadian Institute for Advanced Research (CIFAR), Azrieli Programme on Brain, Mind, and Consciousness.

